# Unlike Chloroquine, mefloquine inhibits SARS-CoV-2 infection in physiologically relevant cells and does not induce viral variants

**DOI:** 10.1101/2021.07.21.451321

**Authors:** Carolina Q. Sacramento, Natalia Fintelman-Rodrigues, Suelen S. G. Dias, Jairo R. Temerozo, Aline de Paula D. Da Silva, Carine S. da Silva, André C. Ferreira, Mayara Mattos, Vinicius C. Soares, Filipe Pereira-Dutra, Milene D. Miranda, Debora F. Barreto-Vieira, Marcos Alexandre N. da Silva, Suzana S. Santos, Mateo Torres, Rajith K R Rajoli, Alberto Paccanaro, Andrew Owen, Dumith Chequer Bou-Habib, Patrícia T. Bozza, Thiago Moreno L. Souza

## Abstract

Repositioning of clinical approved drugs could represent the fastest way to identify therapeutic options during public health emergencies, the majority of drugs explored for repurposing as antivirals for 2019 coronavirus disease (COVID-19) have failed to demonstrate clinical benefit. Without specific antivirals, the severe acute respiratory syndrome coronavirus 2 (SARS-CoV-2) pandemic continues to cause major global mortality. Antimalarial drugs, such as chloroquine (CQ)/hydroxychloroquine (HCQ) and mefloquine have emerged as potential anti-SARS-CoV-2 antivirals. CQ/HCQ entered the Solidarity and RECOVERY clinical trials against COVID-19 and showed lack of efficacy. Importantly, mefloquine is not a 4-aminoquinoline like CQ and HCQ and has been previously repurposed for other respiratory diseases. Unlike the 4-aminoquinolines that accumulate in the high pH of intracellular lysosomes of the lung, the high respiratory tract penetration of mefloquine is driven by its high lipophilicity. While CQ and HCQ exhibit activity in Vero E6 cells, their activity is obviated in TMPRSS2-expressing cells, such as Calu-3 cells, which more accurately recapitulate in vivo entry mechanisms for SARS-CoV-2. Accordingly, here we report the anti-SARS-CoV-2 activity of mefloquine in Calu-3 type II pneumocytes and primary human monocytes. Mefloquine inhibited SARS-CoV-2 replication in Calu-3 cells with low cytotoxicity and EC_50_ and EC_90_ values of 1.2 and 5.3 µM, respectively. In addition, mefloquine reduced up to 68% the SARS-CoV-2 RNA levels in infected monocytes, reducing viral-induced inflammation. Mefloquine blocked early steps of the SARS-CoV-2 replicative cycle and was less prone than CQ to induce drug-associated viral mutations and synergized with RNA polymerase inhibitor. The pharmacological parameters of mefloquine are consistent with its plasma exposure in humans and its tissue-to-plasma predicted coefficient points that this drug may accumulate in the lungs. These data indicate that mefloquine could represent an orally available clinically approved drug option against COVID-19 and should not be neglected on the basis of the failure of CQ and HCQ.

## Introduction

Repurposing of clinically approved drugs represents the fastest way to identify therapeutic options during public health emergencies (1,2). Indeed, the World Health Organization (WHO) and the University of Oxford launched Solidarity and RECOVERY international clinical trials, respectively, against 2019 coronavirus disease (COVID-19) (3,4) just few months after the severe acute respiratory syndrome coronavirus 2 (SARS-CoV-2) emerged in China (5). WHO’s Solidarity trial failed to demonstrate clinical benefit of chloroquine (CQ)/hydroxychloroquine (HCQ), lopinavir (LPV)/ritonavir (RTV) with or without interferon and remdesivir (RDV) (6). LPV/RTV and HCQ were also not effective against COVID-19 in RECOVERY trial (7,8). Conversely, independent clinical investigations on RDV demonstrated its clinical benefit when given early after onset of illness (9–11). Despite that, RDV access is limited to wealthy countries and its intravenous administration is impractical for early and daily use.

Repurposing of immunomodulatory drugs has yielded several successful interventions for later stages of disease (steroids and IL-6 antagonists) (12) and vaccines have been successfully developed at unprecedented speed (13). However, with the slow global roll out of vaccines and without specific antiviral treatments, SARS-CoV-2 continues to cause huge global morbidity and mortality with over 100.000 death/month since its outbreak (14). In addition, new waves of the pandemic continue to cause public health and economic burden globally (15). Libraries of clinically approved compounds have been tested against SARS-CoV-2 (16–18), but consistent antiviral activity in different cellular systems is not common (17). Since severity of COVID-19 is associated with active SARS-CoV-2 replication in type II pneumocytes, cytokine storm and exacerbation of thrombotic pathways (19–21), antiviral compounds identified in Vero cells should be validated in Calu-3 type II pneumocytes and monocytes. Indeed, poor clinical results of CQ/HCQ (6,8) could have been anticipated by the lack of SARS-CoV-2 susceptibility to these drugs in Calu-3 cells and the observation that systemic concentrations did not reach concentration needed to inhibit viral replication.

Very often, antimalarial drugs emerge from compound screening libraries where in vitro activity against SARS-CoV-2 is assessed in unmodified Vero cells (16–18). However, the postulated mechanism of action for CQ and HCQ is via blocking vesicular entry of the virus (22) and these drugs do not retain activity when Vero cells are transfected with TMPRSS2 (23), which represents the predominant entry mechanism in vivo. Unlike CQ and HCQ, mefloquine is not a 4-aminoquinoline, and has been repurposed for other respiratory diseases (e.g. tuberculosis) (24,25) warranting further investigation for SARS-CoV-2. The accumulation of 4-aminoquinolines in the lung is driven by the high pKa of the molecules, which drives lysosomal trapping of the drug in tissues with high lysosome content such as the lung (17). Conversely, high lung concentrations of mefloquine are driven by its lipophilicity giving it has high volume of distribution in lung and other potential infection sites for SARS-CoV-2 (17). These characteristics could be highly beneficial for pulmonary and extra-pulmonary manifestations of COVID-19 (26). Although mefloquine is endowed with activity against other highly pathogenic coronaviruses – SARS-CoV and Middle East respiratory syndrome coronavirus (MERS-CoV) (27) – this drug has been overlooked in clinical trials against COVID-19 (just one study out of more than 5,000 at clinicaltrials.gov). Despite historical concerns about safety of mefloquine, in recent years the safety profile has been positively reviewed (24), for a vulnerable population, pregnant women. The United States Food and Drug Administration (FDA) modified mefloquine safety from class C to B during pregnancy (28).

Robust pre-clinical evaluation should be a prerequisite for progression of candidates into clinical trials. Accordingly, we further characterized the SARS-CoV-2 susceptibility to mefloquine in type II pneumocytes and monocyte cells in which CQ/HCQ failed to produce antiviral activity. Since our previous work indicated that antiviral concentrations of mefloquine may be achievable at approved doses of the drug, we developed a physiologically-based pharmacokinetic model to assess optimal doses for achieving therapeutic concentrations. The presented data strongly indicate that that mefloquine blocks early stages of the virus life cycle and displayed *in vitro* pharmacological parameters consistent with its systemic exposures in humans.

## Materials and Methods

### Reagents

Mefloquine (MQ) and chloroquine (CQ) were received as donations from Instituto de Tecnologia de Fármacos (Farmanguinhos, Fiocruz). Remdesevir (RDV), was purchased from Selleckhem (https://www.selleckchem.com/). The inhibitors were dissolved in 100% dimethylsulfoxide (DMSO) and diluted at least 10^4^-fold in culture medium before each assay. The final DMSO concentrations showed no cytotoxicity. The materials for cell culture were purchased from Thermo Scientific Life Sciences, unless otherwise mentioned. Kits for ELISA assays were purchased from R&D Bioscience.

### Cells and virus

African green monkey kidney (Vero, subtype E6) and human lung epithelial (Calu-3) cell lines were cultured in high glucose DMEM. The culture mediums were complemented with 10% fetal bovine serum (FBS; HyClone), 100 U/mL penicillin and 100 μg/mL streptomycin (P/S) and cells were kept in a humidified atmosphere with 5% CO2 at 37 °C.

Primary monocytes were obtained through the 3h-adherence to plastic from peripheral blood mononuclear cells (PBMCs). PBMCs were isolated from healthy donors’ buffy coat preparations by density gradient centrifugation (Ficoll-Paque, GE Healthcare). Briefly, 2.0 × 10^6^ PBMCs were plated onto 48-well plates (NalgeNunc) in RPMI-1640 without serum. Non-adherent cells were washed and the remaining monocytes were maintained in DMEM high glucose with 5% human serum (HS; Millipore) and P/S. The purity of human monocytes was above 95%, as determined by flow cytometric analysis (FACScan; Becton Dickinson) using anti-CD3 (BD Biosciences) and anti-CD16 (Southern Biotech) monoclonal antibodies.

The SARS-CoV-2 D614G (GenBank #MT710714) and Gamma strains (#EPI_ISL_1060902; kindly donated by Dr. Ester Sabino from Tropical Medicine Institute of the University of São Paulo) were grown in Vero-E6 cells at multiplicity of infection (MOI) of 0.01. Originally, the isolate was obtained from a nasopharyngeal swab from a confirmed case in Rio de Janeiro, Brazil (GenBank #MT710714). All procedures related to virus culture were handled in a biosafety level 3 (BSL3) multiuser facilities according to WHO guidelines. Virus titters were determined as plaque forming units (PFU)/mL. Virus stocks were kept in - 80 °C ultralow freezers.

### Cytotoxicity assay

Monolayers of 2 × 10^4^ cells in 96-well plates were treated with various concentrations of tested drugs (ranging from 1 to 600 μM) for 72 h at 5% CO_2_ and 37 °C. After the incubation period, culture medium was removed, cells were washed with PBS and incubated for 1f at 37°C with a solution containing 1.25% of glutaraldehyde and 0.6 % of methylene blue diluted in Hank’s balanced salt solution (HBSS). Cells were then washed with distilled water, dried at room temperature (RT) and incubated with the elution solution (50 % ethanol, 49 % PBS and 1 % acetic acid) for 1h at RT. The elution solution was transferred to a new 96-well plate and, after a centrifugation at 12,000 x g for 3 minutes, the plate was read at a wavelength of 570 nm (29).

### Yield reduction assay and virus titration

Different cell lineages as Vero-E6 and Calu-3, or primary human monocytes, were plated in 48- or 96-well plates at density of 5 × 10^5^ or 2 × 10^4^ cells/well, respectively, for 24 h at 37 °C. Vero-E6 cells were infected with SARS-CoV-2 at MOI of 0.01. Calu-3 and primary monocytes were infected at MOI of 0.1. After 1 h-incubation period, inoculum was removed and cells were treated with different concentrations of tested drugs diluted in DMEM with 2-10% FBS. After 24 h (for Vero-E6 and primary monocytes) or 48 h (for Calu-3), supernatants were collected for virus titration through PFU/mL or real time RT-PCR and monolayers were lised for quantification of cell-associated viral genomic and subgenomic RNA by real time RT-PCR.

For virus titration, Vero-E6 in 96-well plates (2 × 10^4^ cells/well) were infected with serial dilutions of yield reduction assays’ supernatants containing SARS-CoV-2 for 1h at 37 °C. Semi-solid high glucose DMEM medium containing 2% FSB and 2.4% carboxymethylcellulose (CMC) was added and cultures were incubated for 3 days at 37 °C. Then, the cells were fixed with 3.7% formaline for 2 h at room temperature. The cell monolayer was stained with 0.04% solution of crystal violet in 20% ethanol for 1 h. The virus titers were determined by plaque-forming units (PFU) per milliliter.

### Quantification of viral RNA levels

The total viral RNA from culture supernatants and/or monolayers was extracted using QIAamp Viral RNA (Qiagen®), according to manufacturer’s instructions. Quantitative RT-PCR was performed using GoTaq® Probe qPCR and RT-qPCR Systems (Promega) in an StepOne(tm) Real-Time PCR System (Thermo Fisher Scientific). Primers, probes, and cycling conditions recommended by the Centers for Disease Control and Prevention (CDC) protocol were used to detect the SARS-CoV-2 (30). The standard curve method was employed for virus quantification. For reference to the cell amounts used, the housekeeping gene RNAse P was amplified. Alternatively, genomic (ORF1) and subgenomic (ORFE) were detected, as described elsewhere (31).

### Time of addition and adsorption inhibition assays

Time-of-addition assays were performed in 96-well plates of Calu-3 cells (2 × 10^5^ cells/well). Cells were infected with MOI of 0.1 for 1h at 37 °C. Treatments with mefloquine started from 2 h before to 18 h after infection with two-time EC_50_ concentration. On the next day, culture supernatants were collected and tittered by PFU/mL.

To evaluate the effects of mefloquine on SARS-CoV-2 attachment, monolayers of Calu-3 cells in 96-well plates (2 × 10^5^ cells/well) were infected with MOI of 0.1, for 1h at 37°C. Two different approaches were used: i) SARS-CoV-2 was pre-incubated with 1 μM of mefloquine for 1 h at 37 °C and then used to infect Calu-3 for an additional hour; or ii) Calu-3 cells were pre-treated with 1 μM of mefloquine for 1 h at 37 °C and then infected with SARS-CoV-2. After 24 h, supernatants were collected for virus titration through PFU/mL.

### Measurement of inflammatory mediators

The levels of TNF-α and IL-6 were quantified in the supernatants of SARS-CoV-2-infected monocytes using commercially available ELISA kits, according to manufaturer’s intructions, as we described elsewhere (32).

### Generation of mefloquine-induced mutants

Vero-E6 cells were infected with SARS-CoV-2 at a MOI 0.01 for 1h at 37 °C and then treated with sub-optimal doses of mefloquine or chloroquine. Cells were accompanied daily up to the observation of cytopathic effects (CPE). Virus suspensions were recovered from the culture supernatant, tittered and used in a next round of infection in the presence of higher drug concentrations, ranging ed from 0.5 to 7 µM. As a control, SARS-CoV-2 was also cultured in the absence of treatments to monitor genetic drifts associated with culture passages. Virus RNA was extracted by Qiamp viral RNA (Qiagen) and quantified using Qbit 3 Fluorometer (Thermo Fisher Scientific), according to manufactures’ instructions. The virus RNA was submitted to unbiased sequence using a MGI-2000 and a metatranscriptomics approach, as previously described by us (33). Sequencing data were initially analysed s in the usegalaxy.org platform and then aligned through clustalW, using the Mega 7.0 software.

The susceptibility of emergent mutant isolates to mefloquine and chloroquine was verified in Vero-E6 cells by performing yield reduction assays.

### Transmission Electron Microscopy

Vero-E6 monolayers of 2 × 10^5^ cells in 25 cm^2^ culture flasks were infected with SARS-CoV-2 at a MOI of 1 for 1h at 37 °C. Inoculum was removed and cells were treated with 10 μM of mefloquine. After 4 h, cells were washed with PBS and fixed in 2.5% glutaraldehyde in sodium cacodilate buffer (0.2 M, pH 7.2), post-fixed in 1% buffered osmium tetroxide, dehydrated in acetone, embedded in epoxy resin and polymerized at 60°C over the course of three days (34,35). Ultrathin sections (50– 70 nm) were obtained from the resin blocks. The sections were picked up using copper grids, stained with uranyl acetate and lead citrate (36), and observed using Hitachi HT 7800 transmission electron microscope.

### Differential expression analysis

We performed three different analyses of differential expression. Our first analysis was aimed at identifying pathways targeted by mefloquine. We used data from the Connectivity Map (CMAP) (37,38), a database containing gene expression profiles of cells treated with thousands of compounds. Among the different cell lines available in CMAP, we chose data from A549 cells, which is the one most similar to Calu-3 used in our wet lab experiments. The set of gene comprising the mefloquine expression signature was defined using modified z-scores obtained when comparing mefloquine treated cells with controls across 3 replicates. The data is available from GEO (https://www.ncbi.nlm.nih.gov/geo/) (39), with accession number GSE92742. Finally, to identify pathways targeted by mefloquine, we applied GSEA (40) using KEGG (Kyoto Encyclopedia of Genes and Genomes) pathways (41). GSEA computes an enrichment score that measures how much a pathway is enriched in differentially expressed genes. Pathways with FDR q-value < 0.05 and positive enrichment score were considered to be significantly up-regulated.

Our second analysis was aimed at identifying pathways affected by SARS-CoV-2. We followed a similar procedure, that began by downloading gene expression data from GEO with accession number GSE148729 (42). This consists of RNAseq raw reads obtained from two replicates of SARS-CoV-2 infected Calu-3 cell lines (after 4h of infection) and controls. Differentially expressed gene comprising the SARS-CoV-2 infection expression signature were identified using the R package edgeR (43). We then applied GSEA against the KEGG database. Pathways with FDR q-value < 0.05 and negative enrichment score were considered to be significantly down-regulated.

Our third analysis was aimed at evaluating whether mefloquine is likely to have therapeutic effects against COVID-19 based on its gene expression profiles. Here we used the CMAP pipeline (37,38), comparing the mefloquine expression signature with the SARS-CoV-2 infection expression signature. The pipeline computes the connectivity score (τ), an standardized measure ranging from -100 to 100. This score is obtained by comparing the enrichment score of a given drug to all others in a reference database. A negative τ indicates that the drug and disease have opposite gene expression signatures, suggesting that the drug could revert the effects of the disease (44). A connectivity score below –90 was considered statistically significant

### PBPK model

A whole-body PBPK model was developed in Julia programming language v1.5.3 (in Juno v0.8.4) (45) using packages – DataFrames v0.21.7, DifferentialEquations v6.15.0 and Distributions v0.23.8. The PBPK model consists of various compartments representing various organs and tissues of the human body. Mefloquine physicochemical and drug specific parameters used in the construction of the PBPK model are shown (Table S1). The data from this model is exclusively computer generated, therefore no ethics approval was required for this study.

### Model development

A virtual healthy population of one hundred individuals (50% female) between the ages of 18-75 was used in this study and the characteristics such as weight and height of the individual were randomly generated (using an inbuilt programming functionality) from a population with mean and standard deviation such that every individual is unique (46). Various anthropometric equations were used to compute individual organ and tissue weights/volumes (47) and blood flow rates (48). The tissue to plasma ratios were computed with mefloquine physicochemical characteristics such as pKa, logP and fraction unbound using equations from a previous publication (49). For oral drugs, the small intestine was divided into seven compartments and a simultaneous compartmental absorption and transit (CAT) model was used to simulate the physiological absorption process (50). The drug disposition was defined using various flow equations (51). The PBPK model has a few limitations for this study: 1) entire administered drug is in solution and available for absorption 2) no reabsorption from the large intestine, 3) drug distribution is blood-flow limited, and 4) instant and uniform drug distribution across a tissue/organ.

### Model validation

Mefloquine model was validated against single doses of 250 mg and 1500 mg and also against multiple doses of 250 mg across various clinical studies. The observed data points were digitized using Web-Plot-Digitiser®. The mefloquine PBPK model was assumed to be validated if the difference between the simulated and observed data points computed using the absolute average fold-error is less than two and the mean simulated pharmacokinetic parameters such as AUC, C_max_ and C_trough_ are less than two-fold from the observed values.

### Model simulations

The EC_90_ value from the experimental settings resulted 1200 ng/ml and 2005 ng/ml in Vero-E6 and Calu-3 cell lines, respectively. However, other literature indicated different EC_90_ values in Vero-E6 cells, therefore an average across available studies (2311 ng/ml, Table S2) was used and doses were optimized based on the higher EC_90_ value (Vero-E6 compared to Calu-3). Optimal doses were predicted such that mefloquine trough concentrations were over the average EC_90_ value in Vero-E6 cells (2311 ng/ml) from day 3 to day 7 for at least 90% of the simulated population.

### Statistical analysis

The assays were performed blinded by one professional, codified and then read by another professional. All experiments were carried out at least three independent times, including a minimum of two technical replicates in each assay. The dose-response curves used to calculate EC_50_ and CC_50_ values were generated by variable slope plot from Prism GraphPad software 8.0. The equations to fit the best curve were generated based on R^2^ values ≥ 0.9. Student’s T-test was used to access statistically significant P values <0.05. The statistical analyses specific to each software program used in the bioinformatics analysis are described above.

### Ethics Statement

Experimental procedures involving human cells from healthy donors were performed with samples obtained after written informed consent and were approved by the Institutional Review Board (IRB) of the Oswaldo Cruz Institute/Fiocruz (Rio de Janeiro, RJ, Brazil) under the number 397-07. The National Review Board approved the study protocol (CONEP 30650420.4.1001.0008), and informed consent was obtained from all participants.

## Results

### SARS-CoV-2 replication is inhibited by mefloquine in Calu-3 cells and human primary monocytes

In a recent screening of FDA-approved drugs in SARS-CoV-2-infected Vero cells, mefloquine raised as a potential hit (16,18,52). However, data related to its antiviral effect in physiologically relevant cellular systems for COVID-19 are not available. Therefore, we further evaluated the antiviral effect of mefloquine in Calu-3 type II pneumocytes and primary human monocytes, cells associated with disease severity (19,53) and cytokine storm (19). For comparison, we also evaluated mefloquine effects in SARS-CoV-2-infected Vero-E6 cells.

Calu-3 and Vero-E6 were infected with SARS-CoV-2 at MOI of 0.1 and 0.01, respectively. After 48h (Calu-3) or 24h (Vero-E6) virus replication was assessed in the culture supernants by plaque assay. Mefloquine inhibited SARS-CoV-2 replication in Calu-3 and Vero-E6 in a dose-dependent manner, with EC_50_ values of 1.9 and 0.6 μM, respectively (Table 1 and Figure 1 A-D). Although CQ presented an EC_50_ of 1.1 μM in Vero cells (Table 1 and Figure 1 C-D), it was inactive in Calu-3 cells – which is consistent with previous *in vitro* investigation (54). In contrast, mefloquine reached efficient inhibition of virus replication with optimal inhibitory concentrations ranging from 3.2 to

**Table 1.** Pharmacological parameters of SARS-CoV-2-infected cells in the presence of mefloquine, chloroquine and remdesevir.

| Cell Types | Mefloquine (μM) | | | | Chloroquine (μM) | | | | Remdesivir (μM) | | | |
| --- | --- | --- | --- | --- | --- | --- | --- | --- | --- | --- | --- | --- |
|  | CC<sub>50</sub> | EC<sub>50</sub> | EC<sub>90</sub> | SI | CC<sub>50</sub> | EC<sub>50</sub> | EC<sub>90</sub> | SI | CC<sub>50</sub> | EC<sub>50</sub> | EC<sub>90</sub> | SI |
| Calu-3 | 19.9 ± 0.06 | 1.9 ± 0.6 | 5.3 ± 0.7 | 11 | ND | >10 | >10 | ND | 480 ± 20 | 0.03 ± 0.02 | 0.3 ± 0.04 | 1.6 x10<sup>4</sup> |
| Vero E6 | 9.2 ± 0.3 | 0.6 ± | 3.2 ± 0.3 | 16 | 259 ± 5 | 1.1 ± | 3.5 ± 0.8 | 235 | 512 ± 30 | 1.4 ± 0.3 | 3.3 ± 0.2 | 366 |

SI – Selectivity Index (calculated based on the ratio of the CC<sub>50</sub> and EC<sub>50</sub> values);
ND – Not determined.

**Figure 1.**
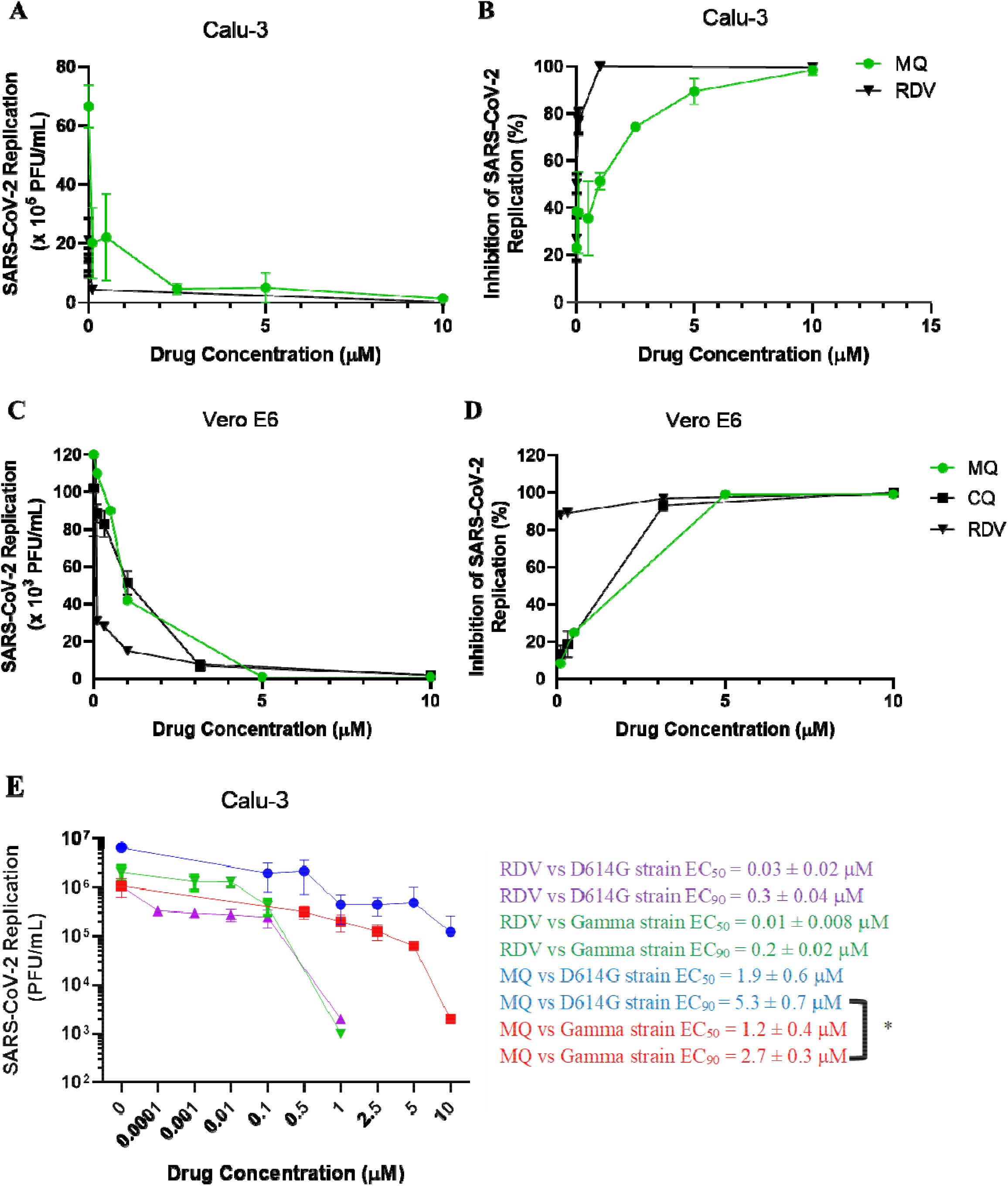
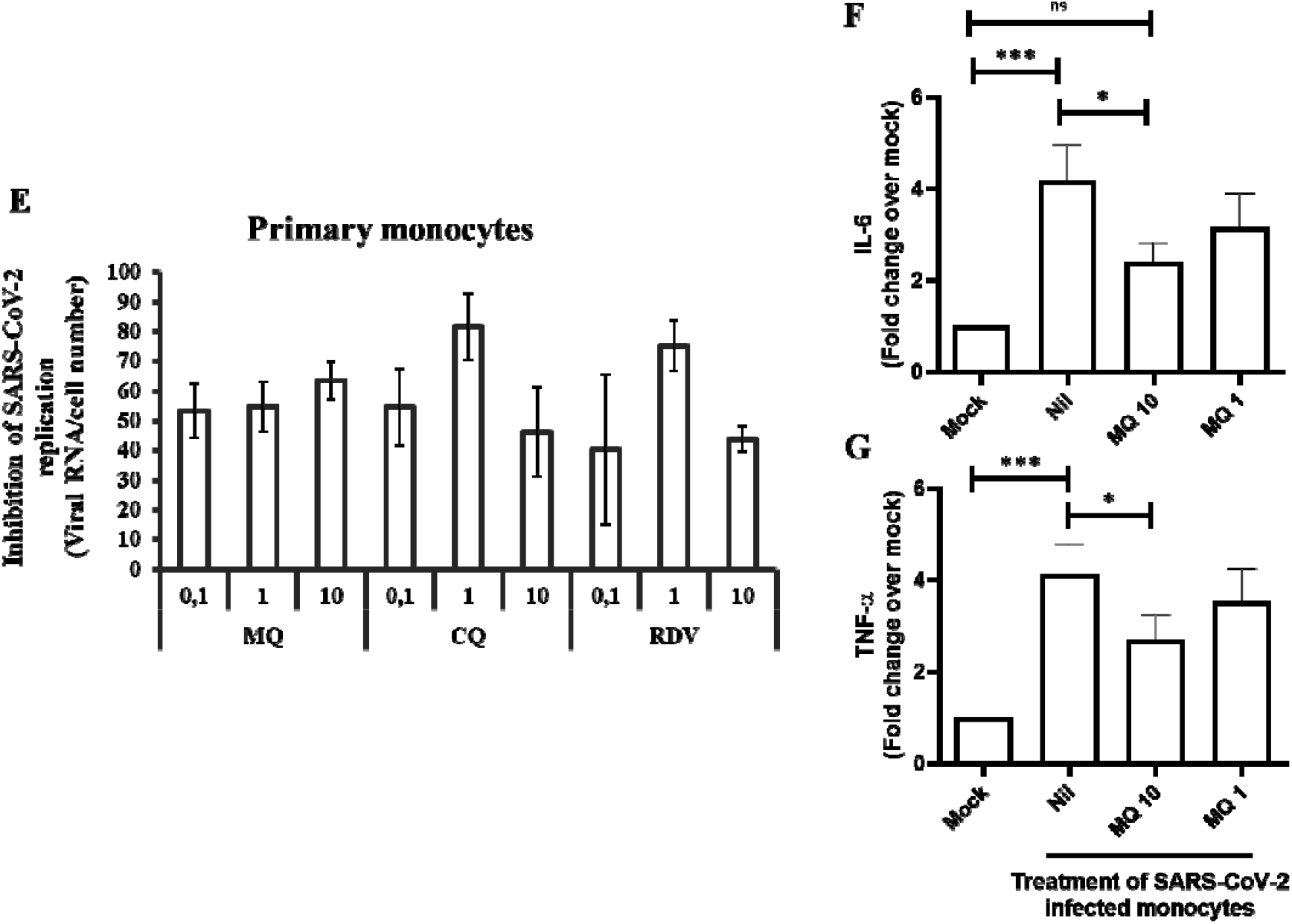
The antiviral effect of mefloquine (MQ) against SARS-CoV-2. Calu-3 (A and B), Vero-E6 (C and D) or primary human monocytes (E to G) were infected with SARS-CoV-2 D614G (A-D and F) or gamma strain (E) at MOI 0.01 (Vero) or 0.1 (Calu-3 and monocytes), for 1h at 37 °C. Inoculum was removed and cells were incubated with fresh DMEM containing the indicated concentrations of mefloquine (MQ), chloroquine (CQ) or remdesevir (RDV). Virus titters were measured by PFU/mL in the culture supernatants after 24 hours post infection (hpi) for Vero (A and B) or 48 hpi for Calu-3 (C and D) cells. Alternatively, viral replication (E), as well as the levels of IL-6 (F) and TNF-α (G), were measured from the monocyte culture supernatants by RT-PCR and ELISA, respectively. Results are displayed as virus titers (A and C) or percentage of inhibition (B, D and E). The data represent means ± SEM of three independent experiments or monocytes from at least 5 healthy donors. *ns* – not significant, \**P*<0.05 and \*\*\**P*<0.01

5.3 μM for Vero-E6 and Calu-3 cells, respectively (Table 1 and Figure 1 A-D). RDV, used as positive control, also displayed suppressive EC_90_ values from 0.3 μM for Vero-E6 and Calu-3 cells, respectively (Table 1 and Figure 1 A-D). The selectivity index (SI), which represents the margin of *in vitro* safety based on CC_50_ and EC_50_ values, highlights that mefloquine exhibits its antiviral activity without cytotoxicity. Moreover, we also evaluated the susceptibility of the variant of concern SARS-CoV-2 gamma to mefloquine in Calu-3 cells and found a slightly higher susceptibility at the efficient level (Figure 1E). As a control, SARS-CoV-2 strains D614G and gamma were similarly susceptible to RDV (Figure 1E).

SARS-CoV-2 can infect immune cells, such as monocytes (55), leading to massive cell death and leukopenia (56). Although human primary monocytes do not produce infectious virus particles, they are susceptible to infection and produce viral RNA (57). Monocytes are key components of the host antiviral response and higher amounts of these cells producing cytokine storm-related mediators are found in patients with severe COVID-19 (53). Mefloquine reduced viral RNA levels in SARS-CoV-2-infected monocytes by 50 to 68%, similarly to CQ and RDV (Figure 1F), as well as the levels of virus-induced monocyte production of IL-6 and TNF-α in infected cultures at 10 µM (Figure 1G and H, respectively).

Altogether, our results indicate that anti-SARS-CoV-2 effect of mefloquine is not restricted to Vero cells, being active also in Calu-3 cells and monocytes.

### Mefloquine is less prone than chloroquine to induce SARS-CoV-2 variants

We next continuously cultured the virus in the presence of mefloquine to monitor emergence of SARS-CoV-2 variants. As a control, virus was also propagated in the presence of CQ or in the absence of antiviral drugs (nil). Following two/three months of successive passages of the virus in Vero-E6 cells at the MOI of 0.1, with increasing concentrations of the drugs, no mefloquine-associated mutant significantly differs from natural virus drifts, because SARS-CoV-2 sequences form mefloquine- or nil (untreated) cells clustered together (Figure 2A, for example see the purple and green circles, which show the highest concentration of mefloquine and its respective control). On the other hand, CQ-associated SARS-CoV-2 variants emerged at 5 µM, differing significantly from their respective control (virus passage in the absence of the drug) (Figure 2A, compare blue and red circles).

**Figure 2.**
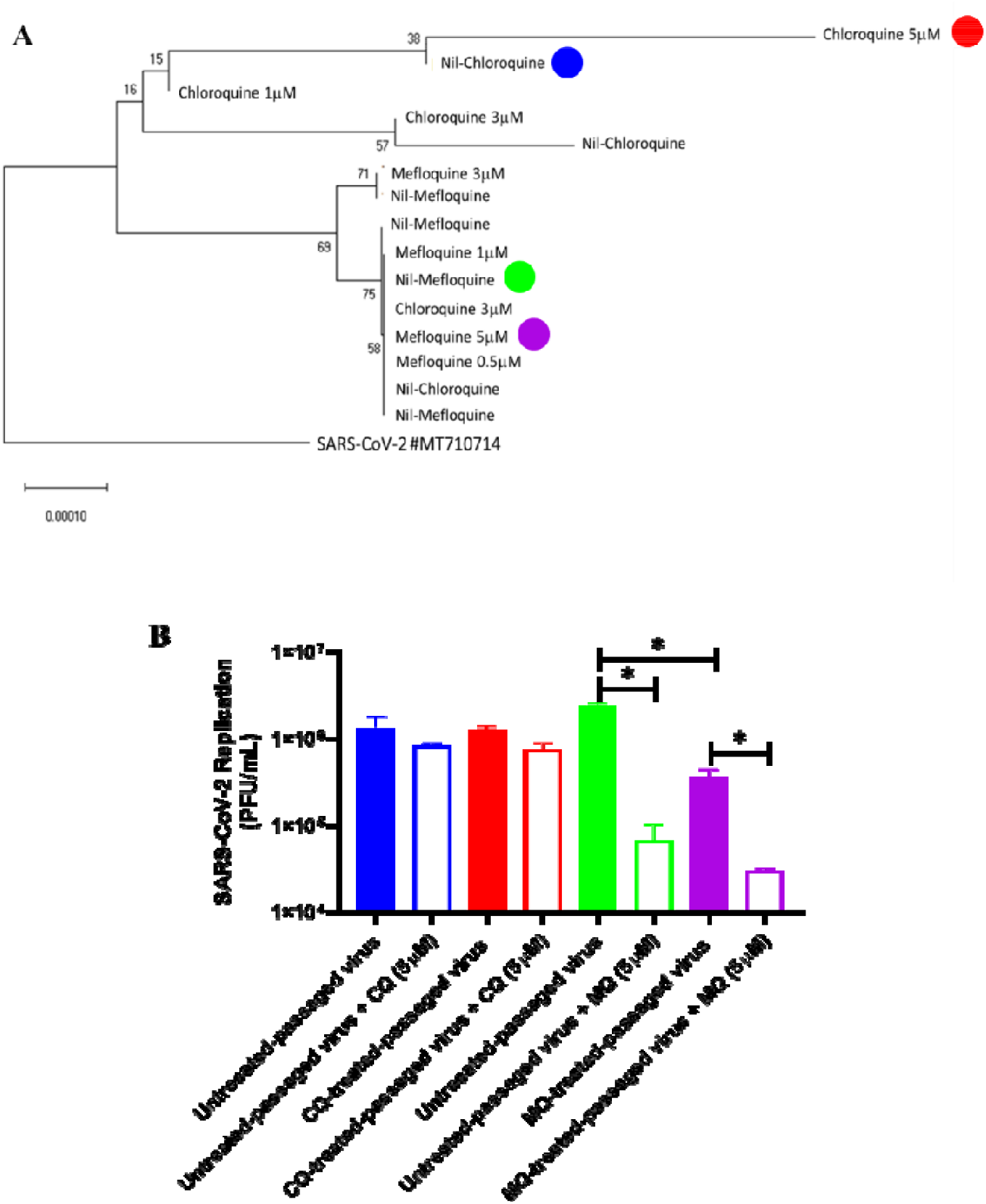
SARS-CoV-2 mutants associated with virus sequencing passage in the presence of mefloquine and chloroquine. (A) SARS-CoV-2 was passaged in Vero-E6 cells, at the MOI of 0.1, in the presence of increasing concentrations of mefloquine or chloroquine (ranging from 0.5 to 7 µM). The virus RNA obtained from the culture supernatants of the passages was submitted to unbiased sequence using MGI-2000 and a metatranscriptomics approach. The virus evolutionary history was inferred using the Neighbor-Joining method (58). The optimal tree with the sum of branch length = 0.00272973 is shown. The percentage of replicate trees in which the associated taxa clustered together in the bootstrap test (1000 replicates) are shown next to the branches (59). The tree is drawn to scale, with branch lengths in the same units as those of the evolutionary distances used to infer the phylogenetic tree. The evolutionary distances were computed using the p-distance method (60) and are in the units of the number of amino acid differences per site. This analysis involved 22 amino acid sequences. The coding data was translated assuming a Standard genetic code table. All ambiguous positions were removed for each sequence pair (pairwise deletion option). There was a total of 9843 positions in the final dataset. Evolutionary analyses were conducted in MEGA X (61). (B) SARS-CoV-2 yielded in the absence or presence of 5 µM of chloroquine (CQ) or mefloquine (MQ) were used to infect Vero-E6 cells (MOI 0.01) for 1h at 37 °C. Inoculum was removed and fresh DMEM containing 5 µM of the drugs was added to the cells. Virus titters were measured by PFU/mL in the culture supernatants after 24 hours post infection (hpi). Results are displayed as virus titers. \**P*<0.05

The virus yielded in either condition, absence or presence of 5 µM of CQ, became naturally resistant to this drug (Figure 2B, compare blue and red bars). After successive passages of SARS-CoV-2 with mefloquine, one-log10-reduction in virus yield indicates that less infectious viruses were produced (Figure 2B, compare green and purple solid bars). When these viruses were tested once more against mefloquine, they remained susceptible to this drug (Figure 2B, compare green or purple solid vs open bars).

### Mefloquine targets endosome-related pathways and its effects on gene expression are opposite to those induced by the SARS-CoV-2 infection

Anti-malarial drugs are endowed with the ability to change endosomal pH to non-physiological (62), such as CQ that blocks virus endocytosis (52,62). To get insights on whether mefloquine would also affect endosome-related pathways, we identified differentially expressed pathways in cell lines treated with mefloquine. Out of 162 KEGG pathways, we found 10 with significant positive enrichment scores (FDR q-value < 0.05), meaning that 10 cellular pathways are up-regulated by mefloquine (Table S3). Among them, six are involved in the endocytosis process (Figure 3).

**Figure 3.**
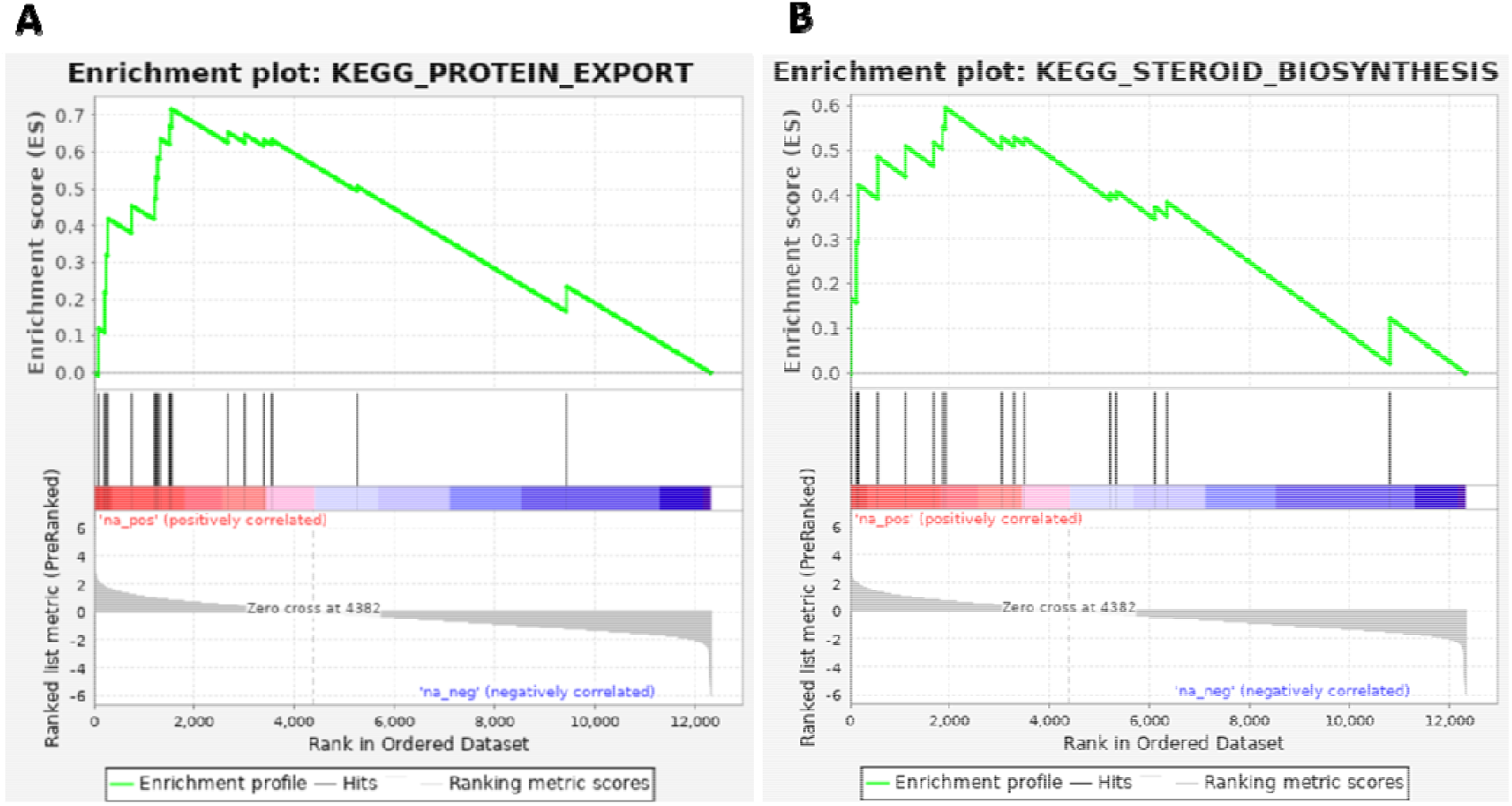
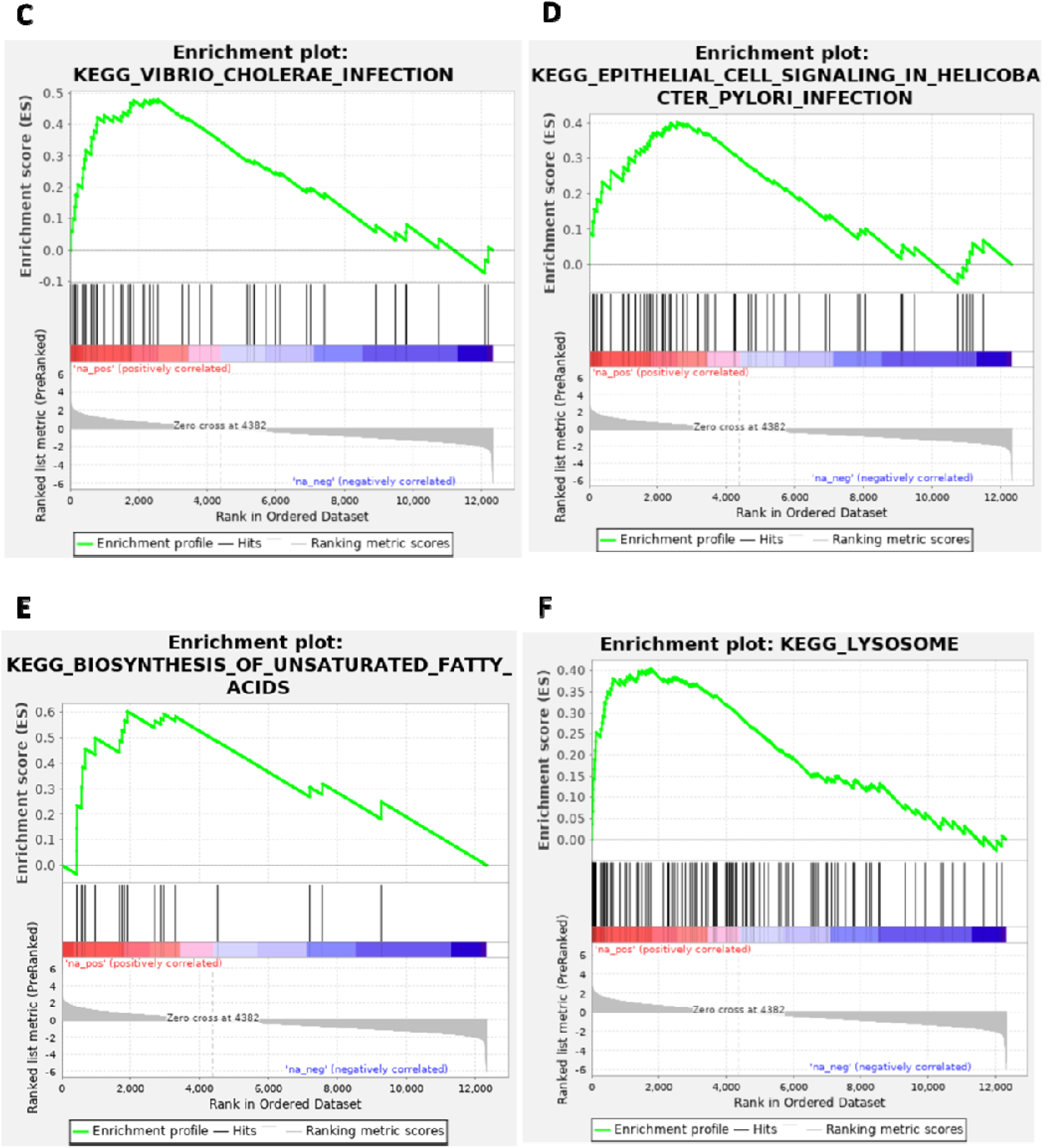
GSEA enrichment plots of the six endocytosis-related pathways up-regulated by mefloquine. We ran GSEA on the mefloquine gene expression signature obtained from CMAP to identify KEGG pathways enriched in differentially expressed genes. The six endocytosis-related pathways with significant enrichment score (FDR < 0.05) are shown here. Subplots contain an identical colored bar, representing the complete gene list ordered by changes in expression levels induced by mefloquine, with red indicating the highest z-scores (up-regulated genes), and blue the lowest ones (down-regulated genes). Each subplot corresponds to a pathway and black vertical lines indicate the position of its genes in the ordered list. Note that for all six pathways, these lines tend to be located towards the top of list (up-regulated genes). The GSEA enrichment score is computed iteratively across the ordered list, as shown in green. The final GSEA enrichment score of each pathway (reported on Table S3) is the highest deviation from zero. The bottom portion of the plot shows the z-score (*y*-axis) at each position of the list (*x*-axis).

Furthermore, to verify whether mefloquine has potential therapeutic effects against COVID-19, we followed an approach first proposed by Sirota *et al*. (44) in which we tested whether the effects of mefloquine on these pathways and on global gene expression are *opposite* to the effects induced by the SARS-CoV-2 infection. First, we identified differentially expressed pathways in Calu-3 cell lines infected by SARS-CoV-2. We found that two out of the six endocytosis-related pathways that are up-regulated by mefloquine are down-regulated by the infection (Table S4), suggesting that mefloquine can oppositely modulate the effect caused by SARS-CoV-2 on these pathways. Then, we ran the CMAP pipeline (37,38) to test whether global mefloquine gene expression signature is opposite to SARS-CoV-2 infection. We found a significant CMAP connectivity score (τ = -90.45), reinforcing the evidence for the potential therapeutic effects of mefloquine.

To confirm that mefloquine impairs SARS-CoV-2 endocytosis-mediated entry, either Calu-3 cells or the virus was pre-treated with mefloquine at 1 µM for 1h. Subsequently, Calu-3 cells were infected with SARS-CoV-2 at MOI of 0.1, and virus production was assessed 24h after infection. In both conditions mefloquine reduced by 3-log_10_ the SARS-CoV-2 replication (Figure 4A). The antiviral effect of mefloquine declined substantially when drug addition was delayed by 4, 6 or 12 hs after infection (Figure 4B). Within 4h after infection, SARS-CoV-2 enters into host cells (Figure 4C and D) and, in fact, mefloquine reduced the number of viral particles in the endosomes (Figure 4E and F). Our results point out that mefloquine is an inhibitor of SARS-CoV-2 entry.

**Figure 4.**
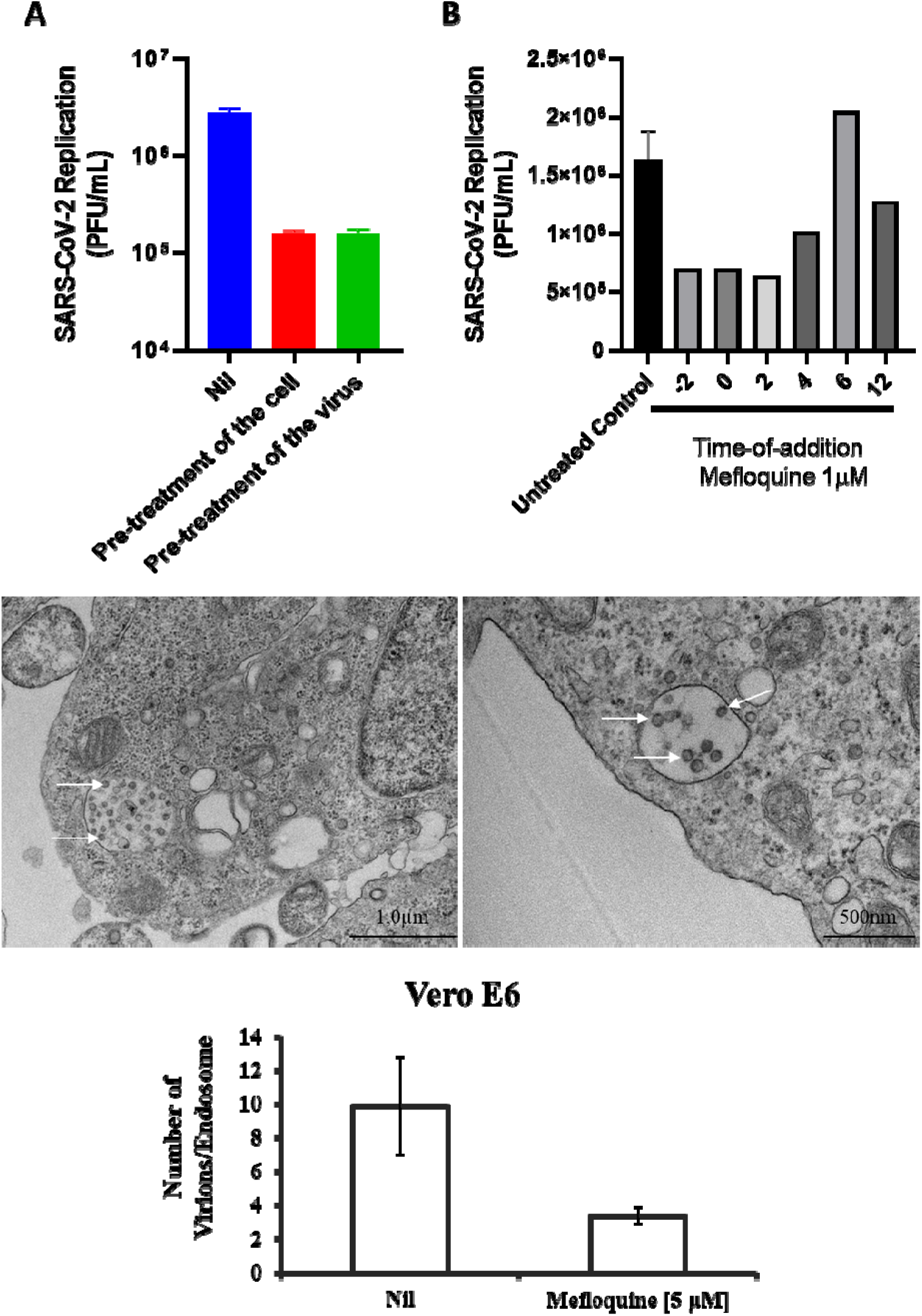
Effects of mefloquine on SARS-CoV-2 endocytosis-mediated entry. (A) To initially evaluate mefloquine’s effect on virus entry, Calu-3 cells or SARS-CoV-2 virus particles were pre-incubated with 1 µM of mefloquine for 1h at 37°C and then infected at MOI of 0.1. (B) Then, a time-of-addition assay was performed in Calu-3 cells infected with SARS-CoV-2 at MOI of 0.1 and treated with 1 µM of mefloquine at different times after infection, as indicated. After 24h post infection, culture supernatants were harvested and SARS-CoV-2 replication was measured by plaque assay. Results are displayed as virus titers. (C-D) Representative images of ultrastructural analysis by transmission electron microscopy of Vero-E6 cell infected with MOI of 1 of SARS-CoV-2 (C) and treated with 5 µM of mefloquine for 4h (D). Cell endosomes with spherical SARS-CoV-2 virus particles (white arrows). (E) SARS-CoV-2 virus particles were counted inside of the endosomes of Vero-E6 cells treated with mefloquine or not (Nil).

Since mefloquine inhibits SARS-CoV-2 entry, it could be expected to synergize with drugs acting on other steps of viral replication. Thus, we combined mefloquine with the currently approved anti-COVID-19 drug remdesivir. RDV’s EC_90_ and EC_99_ were improved by 200-fold in the presence of suboptimal concentrations of mefloquine to inhibit from 25 to 50 % of viral replication (Figure 5). These findings indicate that mefloquine could be accessory drug to improve RDV’s activity, because its short half-life in humans may compromise its robust preclinical results.

**Figure 5.**
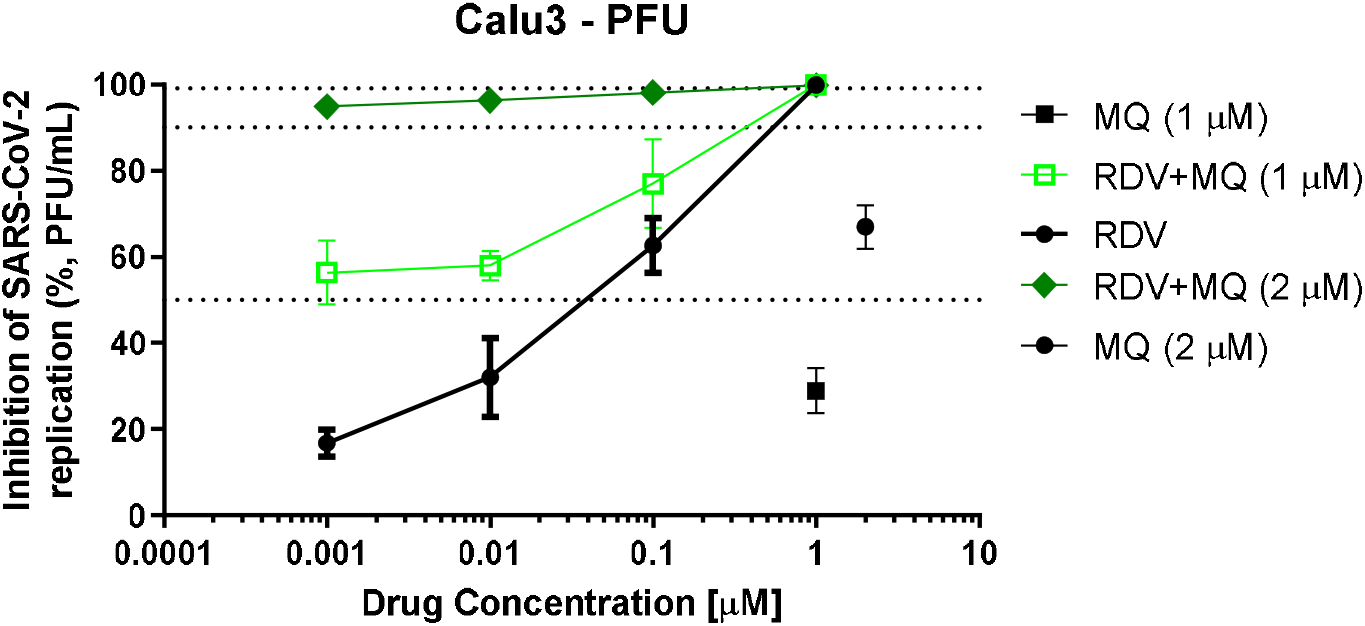
Mefloquine enhances the anti-SARS-CoV-2 activity of remdesivir. Calu-3 cells were infected with SARS-CoV-2 at MOI 0.1 for 1h at 37 °C. Inoculum was removed and cells were treated with 1 or 2 µM of mefloquine combined with the indicated concentrations of remdesivir (RDV). Virus titters were measured by PFU/mL in the culture supernatants after 48 hours post infection (hpi). Results are displayed as percentage of inhibition.

### Mefloquine PBPK modeling and dose simulations

A mefloquine PBPK model was validated against available clinical data and the AAFE of the simulated plots were between 0.96 – 1.21. Figures S1-4 show the comparison between observed and simulated data for various dosing strategies. A scaling factor for intrinsic clearance and tissue plasma ratios of 2 and 0.38 respectively were applied to validate the model. Furthermore, the bioavailability was increased by 10% from day 7 onwards to match the clinical data for multiple dosing studies as mefloquine seems to exhibit higher bioavailability for successive doses.

Various dose settings were assessed and the best two doses i.e. 450 mg TID and 350 mg QID for 3 days are shown in Figures 6A and B such that the plasma concentrations of mefloquine are over the target EC_90_ value in 90% of the simulated population from day 3 to 7. However, much higher doses of mefloquine (1400 mg TID or 1100 mg QID on day 1 only, data not shown) would be required if the plasma concentrations had to reach the target EC_90_ on day 1 and may not be practically possible to administer in individuals. Figures S5 and S6 show the median C_trough_ on day 1, 3 and 7 and the percentage of population over the target EC_90_ value of 2311 ng/ml (or 8.3 µM) for TID and QID settings across various doses.

**Figure 6.**
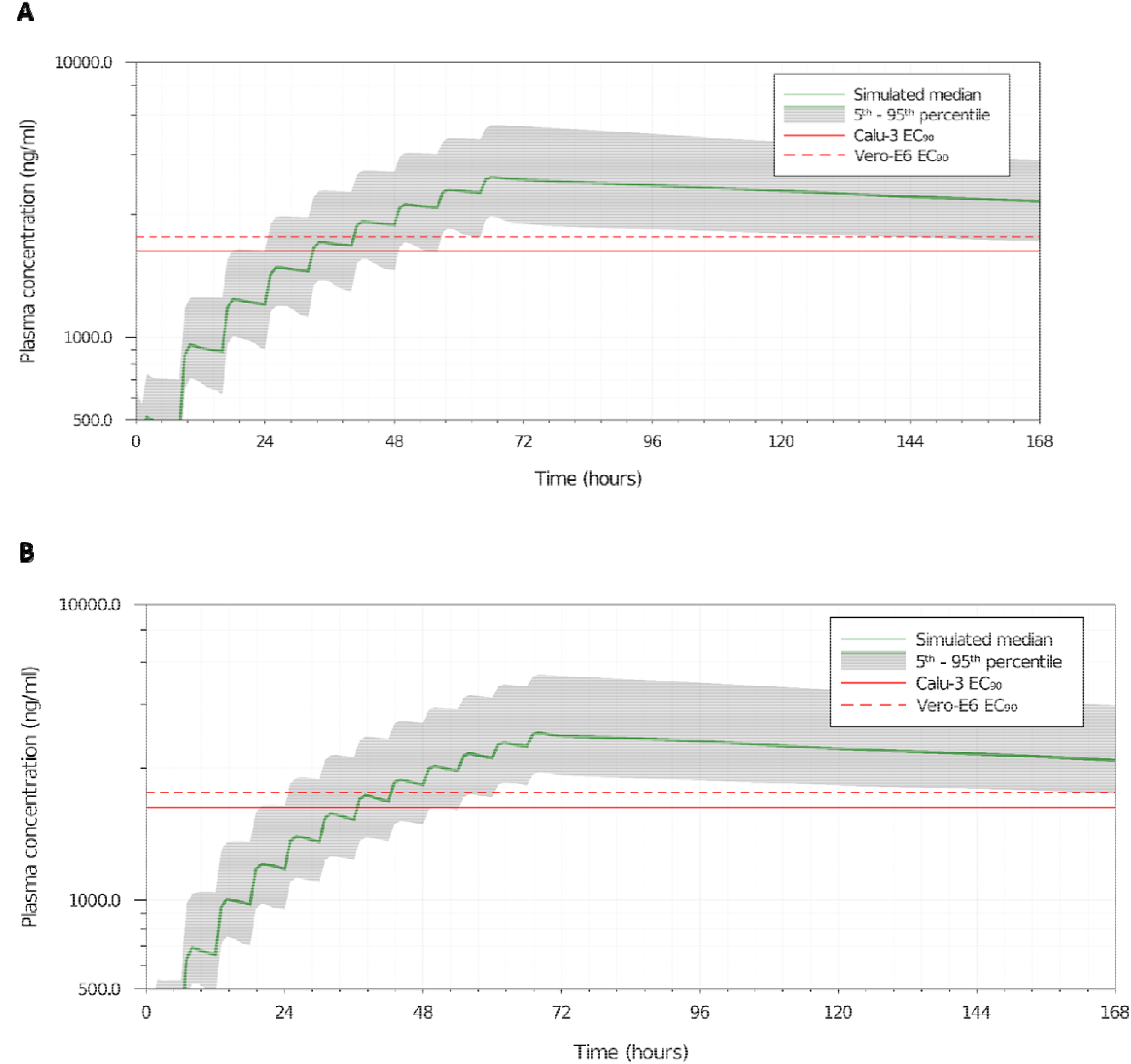
Mefloquine PBPK model predicted plasma concentration for different dosages. (A). Predicted mefloquine plasma concentration for 450 mg TID (B) or 350 mg QID (C) doses for 3 days. The dotted and the solid lines represent the EC_90_ values of mefloquine for SARS-CoV-2 in Vero-E6 and Calu-3 cell types, respectively.

Although the plasma exposure to reach the EC_90_ would require a new therapeutic dosage and regimen, the EC_50_ (756 ng/ml or 2 µM) and EC_25_ (378 ng/ml or 1 µM) are achievable after a single dose of mefloquine at 1500 mg for over 7 days (Figure S2) and has been demonstrated to enhance RDV’s activity.

## Discussion

The clinical evolution of mild to severe COVID-19 is characterized by a multi-stage disease, in which acute viral replication diminishes after the second to third weeks after the onset of illness. In severe COVID-19, patients that required oxygen support present an intense pro-inflammatory response along with coagulopathy (19,63,64). Antivirals are expected to be used on COVID-19 patients to prevent evolution to poor clinical outcomes. RDV, which is clinically approved for COVID-19, showed clinical benefits in Wang *et al* and Beigel *et al* studies (9,11) but failed to do so in other investigations (6). In fact, the poor oral bioavailability and prerequisite for intravenous administration, mitigates widespread roll out in community intervention programs. Orally available drugs endowed with the ability to reduce SARS-CoV-2 levels could reduce hospitalization, the chain of transmission and mortality. Our results show that mefloquine inhibited SARS-CoV-2 replication by blocking virus entry, decreasing anti-inflammatory activity and enhancing RDV’s activity with concentrations close to those achievable with approved doses.

Host-directed broad spectrum antimicrobial drugs have been attempted against COVID-19, such as CQ/HCQ and nitazoxanide (6,65). Whereas *in vitro* data and very first clinical studies on nitazoxanide are under development (65,66), initial enthusiasm with CQ/HCQ is not sustained scientifically (6,8). The clinical failure of CQ/HCQ to enhance recovery of COVID-19 patients could have been anticipated by *in vitro* experimentation and assessment of pharmacokinetic performance (6,54). CQ/HCQ lacks antiviral activity on SARS-CoV-2-infected Calu-3 cells (54) which express TMPRSS2. While the effect of 4-aminoquinolines on endosomal pH could have utility for viruses that depend on this organelle (52,62), the lack of activity in SARS-CoV-2 is now clear.

Amongst antimalarial agents, mefloquine displays good pharmacological properties to be repurposed against COVID-19. Unlike 4-aminoquinolines, mefloquine inhibited SARS-CoV-2 entry in Calu-3 cells, a model for type II pneumocytes; and monocytes, the most affected cells in the respiratory tract of patients that deceased from COVID-19 (19,53). Mefloquine tissue-to-plasma predicted coefficient points that this drug may accumulate in the lungs (17), predominantly via high lipophilicity and not pKa like 4-aminoquinolines. Indeed, mefloquine has been repurposed for tuberculosis, another disease of the respiratory tract (67). Importantly, the *in vitro* pharmacological parameters described here for mefloquine are at least 5-fold better when compared to tuberculosis, which requires a minimal inhibitory concentration of 20 µM (68). Another advantage of mefloquine was the absence of SARS-CoV-2 mutants with decreased sensitivity to this drug, virus particles remained genetically stable upon treatment with mefloquine.

Due to its high lipophilicity, expressed in the form of a logP of 4.0, mefloquine is well distributed throughout anatomical compartments and has a long half-life, measured in weeks (69), allowing that a single or few oral doses could be combined with other antivirals. With C_max_ of 3279 ng/ml, which is equivalent to 8.67 µM (70), and our *in vitro* pharmacological parameters ranging from 0.6 to 5.4 µM, the antiviral activity described here may be physiologically achievable. For comparison, CQ’s C_max_ is 870 ng/ml (or 2.73 µM) (71). Because EC_90_ values for CQ in Vero-E6 cells are higher than its C_max_, it is unfeasible the perspective of its clinical benefit for COVID-19 patients. On the other hand, human plasmatic exposure to mefloquine’s EC_90_ could be achievable under new therapeutic regimens. Mefloquine does exhibit high binding to plasma proteins (>98%) and this adds a degree of uncertainty. However, proteins are included in the *in vitro* activity assays, and the importance of protein binding depends upon multiple drug- and target-specific principles (72). Moreover, even under the current therapeutic approach, suboptimal doses of mefloquine could enhance the anti-SARS-CoV-2 activity of drugs acting on different steps of virus life cycle, which for instance was demonstrated for RDV, but could be likely applicable to other RNA polymerase inhibitors, such as molnupinavir and favipiravir. In addition, short half-life of and the accumulation of prodrug-nucleotides (proTides) in the liver may limit the adequate bioavailability of RDV in the respiratory tract (73). Thus, the combination of mefloquine and RDV could be beneficial. Indeed, mefloquine alters the adenine and purine metabolism (74), which could explain its beneficial effect when combined with RDV.

Since mefloquine toxicological profile was recently reviewed to become a class B product for pregnancy, meaning that it is safer for vulnerable persons, it could represent an orally available clinically approved drug to be considered for clinical trials against COVID-19.

## Acknowledgments

Thanks are due to Dr. Carmen Beatriz Wagner Giacoia Gripp from Oswaldo Cruz Institute for assessments related to BSL3 facility. Dr. Andre Sampaio from Farmanguinhos, platform RPT11M, is acknowledged for kindly donate the Calu-3. Our thanks are also due to Dr. Ester Sabino from Tropical Medicine Institute of the University of São Paulo for kindly donate SARS-CoV-2 Gamma strain. We thank the Hemotherapy Service of the Hospital Clementino Fraga Filho (Federal University of Rio de Janeiro, Brazil) for providing buffy-coats. This work was supported by Conselho Nacional de Desenvolvimento Científico e Tecnológico (CNPq), Fundação de Amparo à Pesquisa do Estado do Rio de Janeiro (FAPERJ). This study was financed in part by the Coordenac□ão de Aperfeic□oamento de Pessoal de Nível Superior - Brasil (CAPES) - Finance Code 001. Funding was also provided by CNPq, CAPES and FAPERJ through the National Institutes of Science and Technology Program (INCT) to Carlos Morel (INCT-IDPN). Thanks are due to Oswaldo Cruz Foundation/FIOCRUZ under the auspicious of Inova program (B3-Bovespa funding). The funding sponsors had no role in the design of the study; in the collection, analyses, or interpretation of data; in the writing of the manuscript, and in the decision to publish the results. A.O. acknowledges funding from EPSRC, which was repurposed from EP/R024804/1 as part of the UK emergency response to COVID-19.

## Notes

### Competing Interest Statement

The authors have declared no competing interest.

