## Supplementary Material for "Unlike Chloroquine, mefloquine inhibits SARS-CoV-2 infection in physiologically relevant cells and does not induce viral variants"

**Table S1.** Mefloquine PBPK input parameters.

| **Parameter** | **Mefloquine** |
| --- | --- |
| Molecular weight | 378.3 [11] |
| Protein binding | 98% [12] |
| Log P | 3.87 [12] |
| pKa (diprotic base) | 8.7 [12] |
| Blood-to-plasma ratio | 1.5 [12] |
| Effective permeability | 6.48 × 10^-4^ cm/s [12] |
| Intrinsic clearance | 10.47 L/h [12] |
| Volume of distribution | 10.2 L/kg^*^ |
| Half-life | 2-4 weeks [11] |

^*^Computed using Rodgers, Leahy and Rowland equation (1).

**Table S2.** Mefloquine average EC_90_ values from various literature sources and experimental settings.

| **Cell type** | **EC_90_ (ng/ml)** | **Reference** |
| --- | --- | --- |
| Vero-E6 | 2513.8 | [10] |
| Vero-E6 | 2455.2 | [9] |
| Vero-E6 | 3064.3 | [8] |
| Vero-E6 | 1210.6 | experimental |
| Average (Vero-E6) | 2311.0 |  |
| Calu-3 | 2005.1 | experimental |

**Table S3.** KEGG pathways significantly up-regulated by mefloquine’s treatment. We ran GSEA enrichment analysis with mefloquine gene expression signature from CMAP. Out of 162 KEGG pathways, ten were found to be up-regulated (FDR < 0.05). For each of these, we report the number of genes involved in the pathway that are on CMAP dataset (column “size”). In addition, we report the GSEA enrichment score (ES), the normalized enrichment score (NES), the nominal p-value testing whether ES is larger than zero (NOM p-val), and the adjusted p-value by the false detection rate (FDR q-val).

| **NAME** | **SIZE** | **ES** | **NES** | **NOM p-val** | **FDR q-val** |
| --- | --- | --- | --- | --- | --- |
| KEGG_PROTEIN_EXPORT | 17 | 0.716 | 2.491 | 0.000 | 0.000 |
| KEGG_VIBRIO_CHOLERAE_INFECTION | 48 | 0.479 | 2.239 | 0.000 | 0.005 |
| KEGG_BIOSYNTHESIS_OF_UNSATURATED_FATTY_ACIDS | 19 | 0.604 | 2.191 | 0.005 | 0.004 |
| KEGG_LYSOSOME | 109 | 0.405 | 2.142 | 0.000 | 0.004 |
| KEGG_STEROID_BIOSYNTHESIS | 16 | 0.595 | 2.060 | 0.000 | 0.005 |
| KEGG_EPITHELIAL_CELL_SIGNALING_IN_HELICOBACTER_PYLORI_INFECTION | 60 | 0.404 | 2.001 | 0.000 | 0.008 |
| KEGG_SPLICEOSOME | 106 | 0.346 | 1.915 | 0.000 | 0.013 |
| KEGG_PROPANOATE_METABOLISM | 27 | 0.471 | 1.852 | 0.010 | 0.019 |
| KEGG_RIBOSOME | 84 | 0.341 | 1.807 | 0.000 | 0.023 |
| KEGG_INOSITOL_PHOSPHATE_METABOLISM | 49 | 0.362 | 1.688 | 0.007 | 0.044 |

**Table S4.** KEGG endocytosis-related pathways down-regulated by SARS-CoV-2 infection. We ran GSEA enrichment analysis with gene expression signature of SARS-CoV-2 infection in Calu-3 cell lines. Out of six endocytosis-related pathways up-regulated by mefloquine, two are significantly down-regulated by SARS-CoV-2 infection. For each pathway, we report the number of corresponding genes available on the expression dataset (size), the GSEA enrichment score (ES), the normalized enrichment score (NES), the nominal p-value testing if ES is smaller than zero (NOM p-val), and the adjusted p-value by the false detection rate (FDR q-val).

| NAME | SIZE | ES | NES | NOM p-val | FDR q-val |
| --- | --- | --- | --- | --- | --- |
| KEGG_PROTEIN_EXPORT | 24 | -0.638 | -1.951 | 0.002 | 0.002 |
| KEGG_STEROID_BIOSYNTHESIS | 15 | -0.623 | -1.674 | 0.010 | 0.026 |

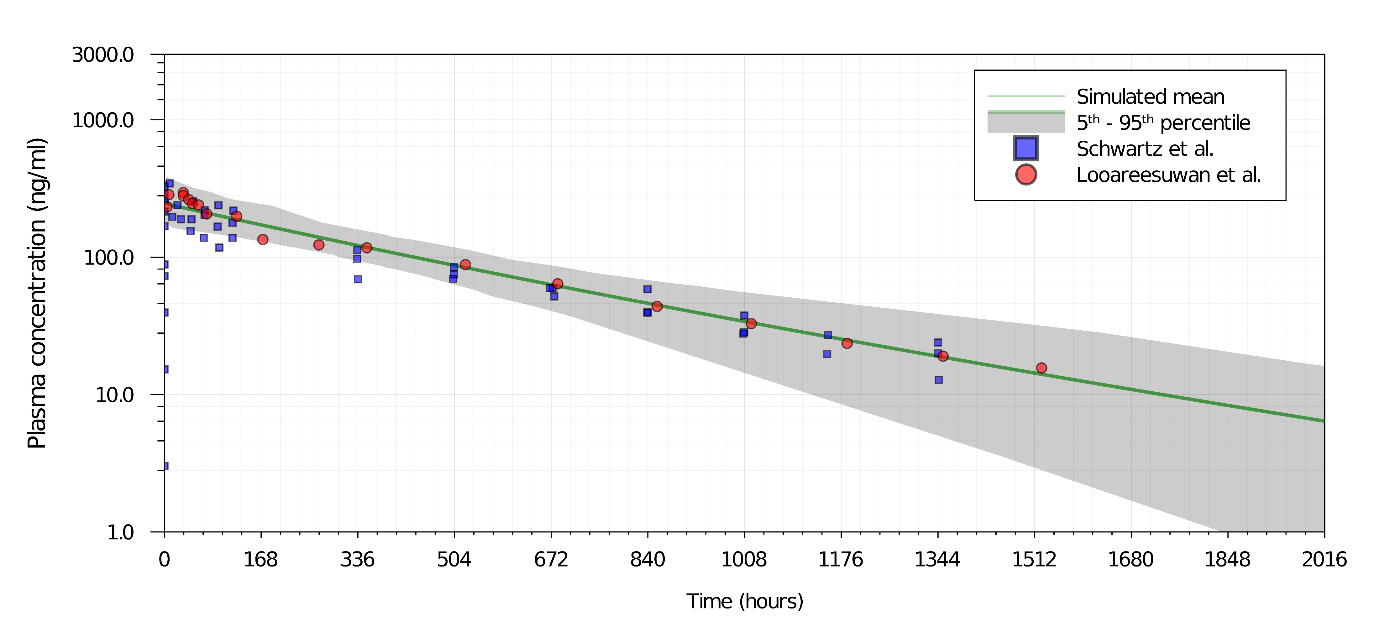

**Figure S1.** Mefloquine PBPK model validation against clinical data (2,3) for a single 250 mg dose.

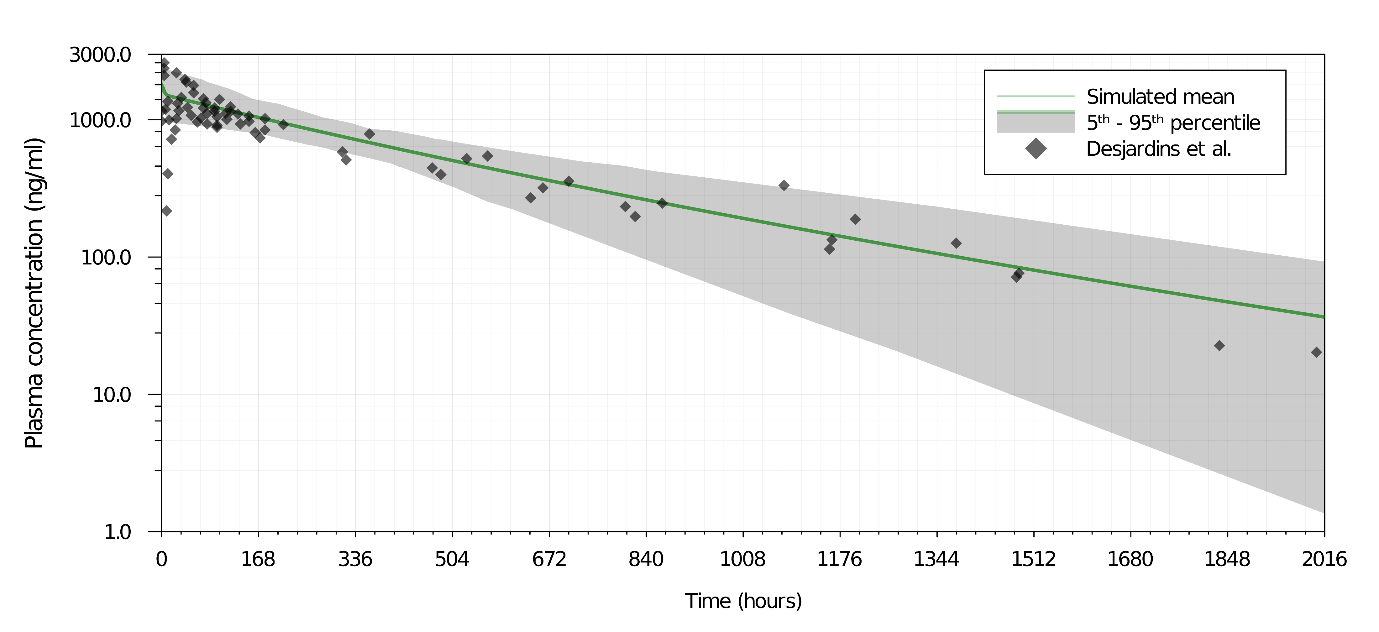

**Figure S2.** Mefloquine PBPK model validation against clinical data (4) for a single 1500 mg dose.

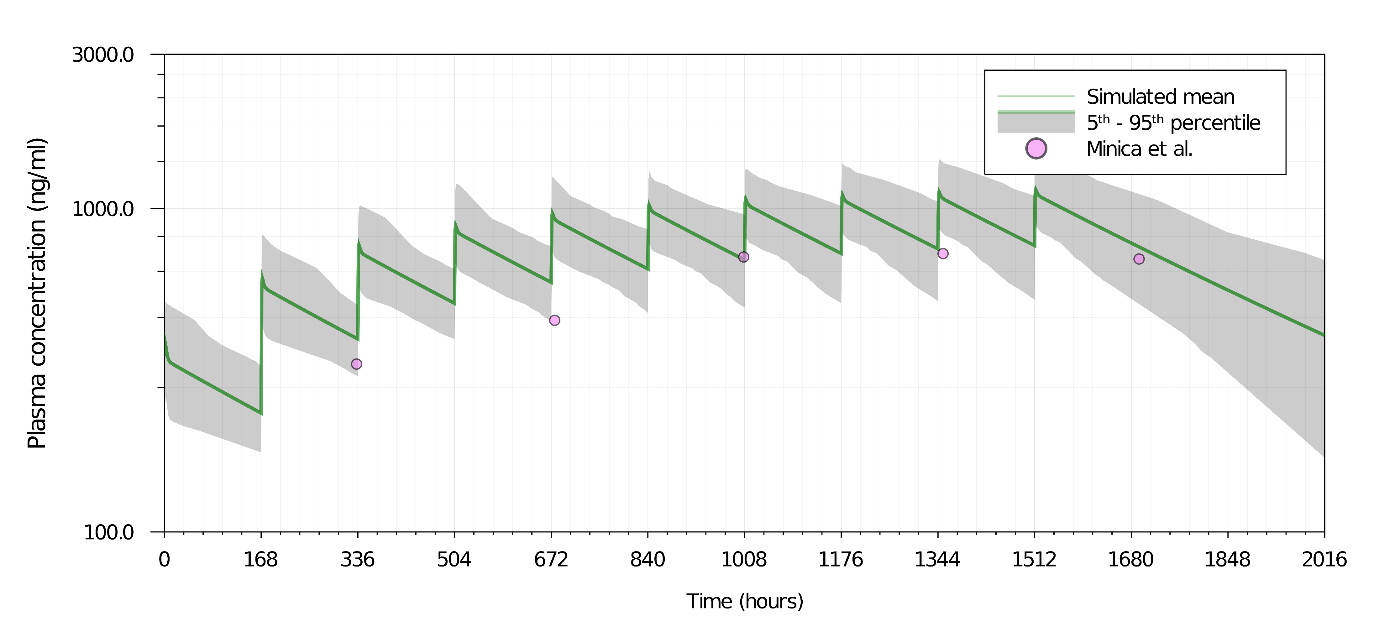

**Figure S3.** Mefloquine PBPK model validation against clinical data (5) for 250 mg doses administered once every week for 10 weeks.

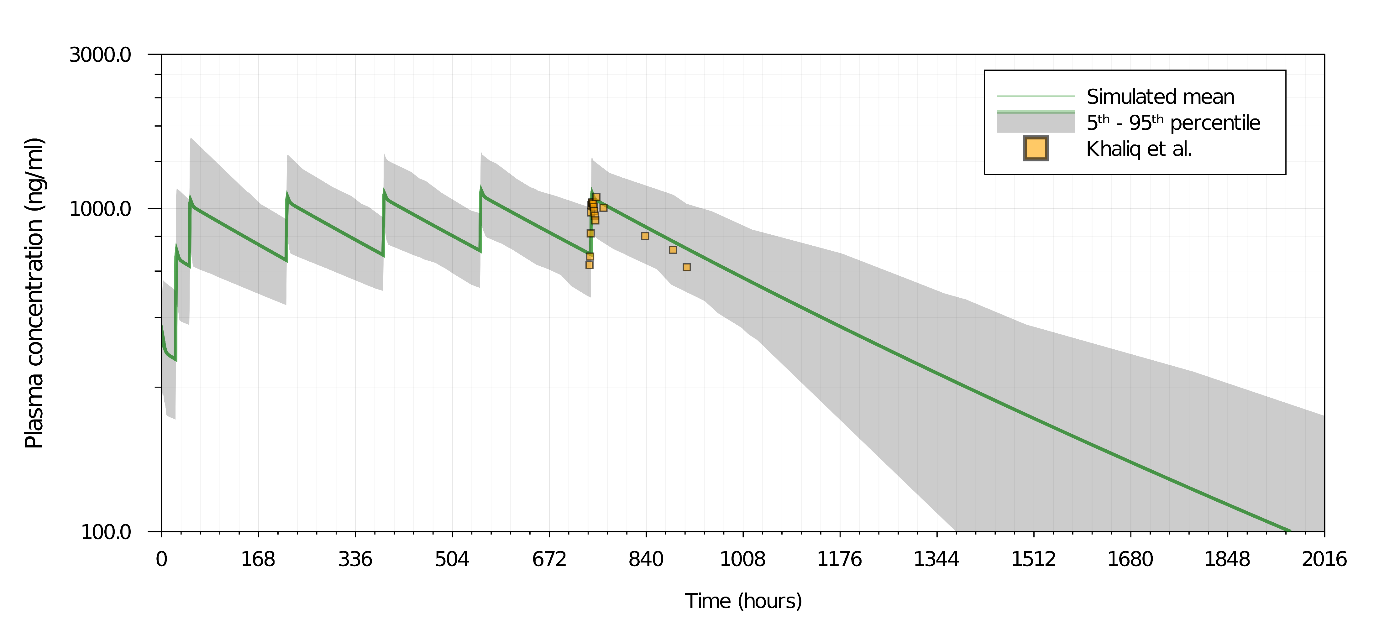

**Figure S4.** Mefloquine PBPK model validation against clinical data (6) for 250 mg doses administered once on day 0, 1, 2 followed by once weekly for 5 weeks.

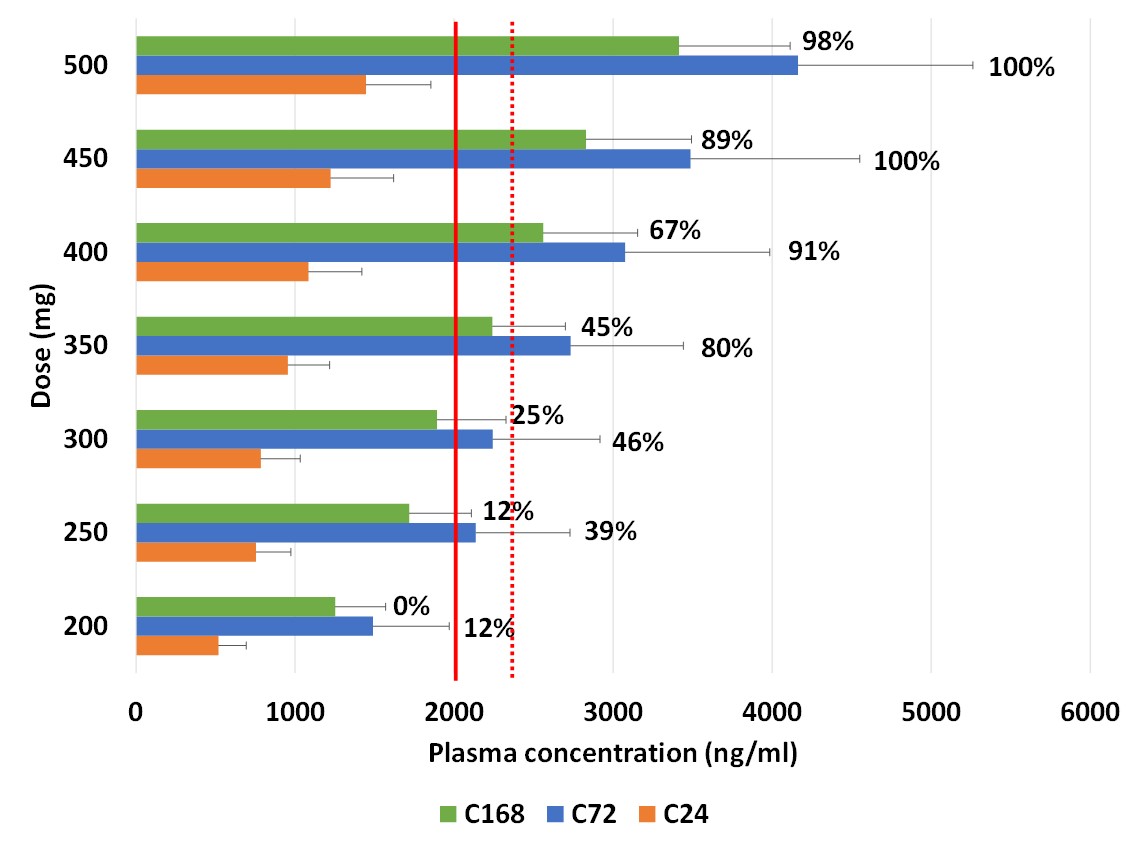

**Figure S5.** Median plasma trough concentrations of mefloquine on day 1 (C24), 3 (C72) and 7 (C168) for various doses administered TID for 3 days. The solid and the dotted red lines represents the EC_90_ values of mefloquine in Calu-3 and Vero-E6 cells for SARS-CoV-2, respectively. The numbers adjacent to each of the bars indicate the percentage of simulated population having plasma concentrations over the target EC_90_ value in Vero-E6 cells

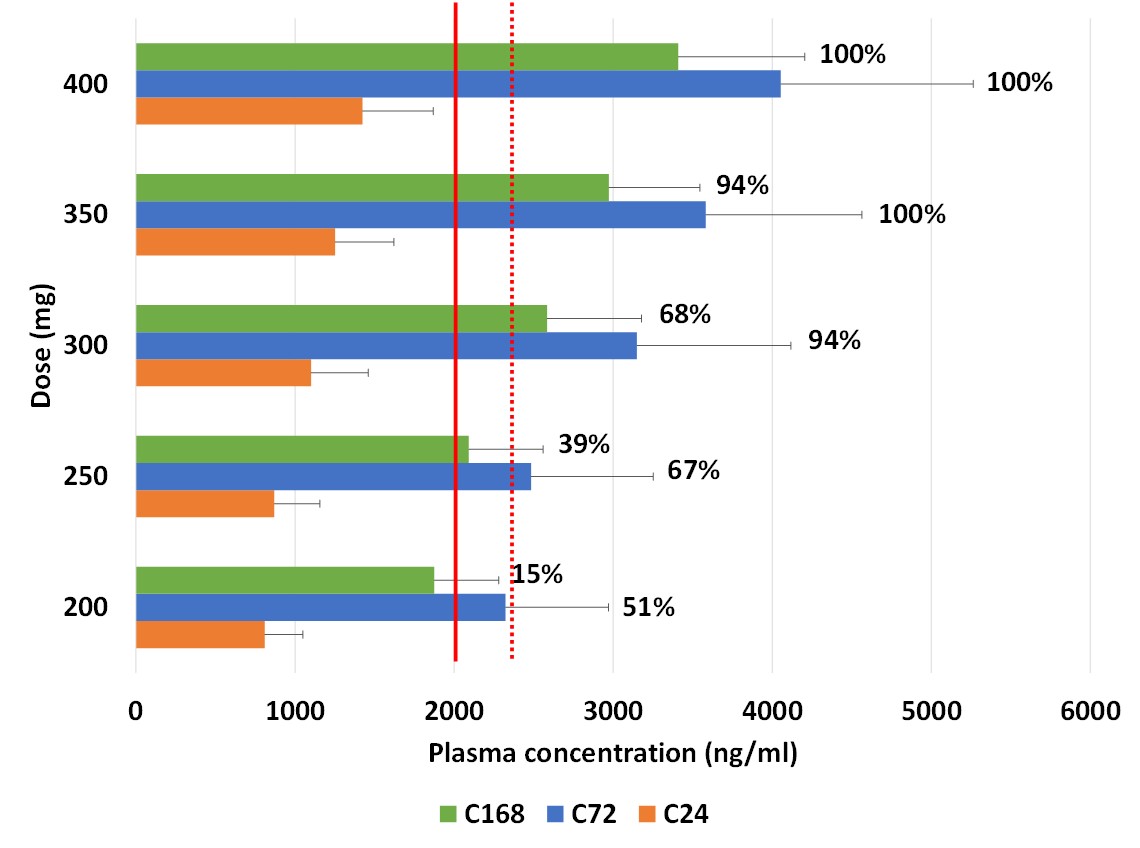

**Figure S6.** Median plasma trough concentrations of mefloquine on day 1 (C24), 3 (C72) and 7 (C168) for various doses administered QID for 3 days. The solid and the dotted red lines represents the EC_90_ values of mefloquine in Calu-3 and Vero-E6 cells for SARS-CoV-2, respectively. The numbers adjacent to each of the bars indicate the percentage of simulated population having plasma concentrations over the target EC_90_ value in Vero-E6 cells.
